# Vaccine-induced protection from homologous Tier 2 simian-human immunodeficiency virus challenge in nonhuman primates

**DOI:** 10.1101/454769

**Authors:** Matthias G. Pauthner, Joseph P. Nkolola, Colin Havenar-Daughton, Ben Murrell, Samantha M. Reiss, Raiza Bastidas, Jérémie Prévost, Rebecca Nedellec, Benjamin von Bredow, Peter Abbink, Christopher A. Cottrell, Daniel W. Kulp, Talar Tokatlian, Bartek Nogal, Matteo Bianchi, Hui Li, Jeong Hyun Lee, Salvatore T. Butera, David T. Evans, Lars Hangartner, Andrés Finzi, Ian A. Wilson, Rich T. Wyatt, Darrell J. Irvine, William R. Schief, Andrew B. Ward, Rogier W. Sanders, Shane Crotty, George M. Shaw, Dan H. Barouch, Dennis R. Burton

## Abstract

Passive administration of HIV neutralizing antibodies (nAbs) can protect macaques from hard-to-neutralize (Tier 2) chimeric simian-human immunodeficiency virus (SHIV) challenge. However, conditions for nAb-mediated protection following vaccination have not been established. Here, we selected groups of 6 rhesus macaques with either high or low serum nAb titers from a total of 78 animals immunized with recombinant native-like (SOSIP) Env trimers from the BG_505_ HIV isolate. Repeat intrarectal challenge with homologous Tier 2 SHIVBG_505_ led to rapid infection in unimmunized and low-titer animals. In contrast, high-titer animals demonstrated protection that was gradually lost as nAb titers waned over weeks to months. From these results, we determined that an autologous serum ID_50_ nAb titer of ~1:500 was required to afford over 90% protection from medium-dose SHIV infection. We further identified autologous nAb titers, but not ADCC or T cell activity, as strong correlates of protection. These results provide proof-of-concept that Env protein-based vaccination strategies can protect against hard-to-neutralize SHIV challenge in rhesus macaques by inducing Tier 2 nAbs, provided appropriate neutralizing titers can be reached and maintained.

## INTRODUCTION

Several vaccine strategies are being pursued to stimulate protective immunity against HIV, including those that combine the elicitation of cellular and humoral responses (Haynes and Burton, 2017; Stephenson et al., 2016). One of the most intensively studied approaches is focused on inducing neutralizing antibodies (nAbs) to the virus. Early pioneering monkey studies showed that DNA/gp120-immunization induces nAb responses that can protect against Tier 1 virus challenge (Barnett et al., 2008; 2010; Pal et al., 2006). However, Tier 1 viruses like SHIV_Ba-L_ and SHIV_SF162-P4_ are easy to neutralize, typically lead to self-limiting infections and are not considered representative of circulating viruses in the HIV pandemic. Two recent studies investigated vaccine-induced protection from a mixed Tier SIVsmE660 swarm and attributed protection, in part, to nAb and other Ab responses (Keele et al., 2017; Roederer et al., 2015). Currently there has not been clear evidence of vaccination-induced nAbs providing protection against viruses possessing hard-to-neutralize clinically relevant Tier 2 HIV Env in humans or NHP models.

Enthusiasm for the nAb approach arises from the association of nAbs with protection for other viruses (Tomaras and Plotkin, 2017) and the demonstration that passively administered HIV-neutralizing monoclonal antibodies (mAbs) can afford protection in monkey and mouse models of HIV infection (Gautam et al., 2016; Gruell et al., 2013; Hessell et al., 2007; Mascola et al., 2000; Moldt et al., 2012; Parren et al., 2001; Pegu et al., 2014; Shingai et al., 2014). As HIV does not infect monkeys, HIV-neutralizing mAbs are assessed by their ability to protect against chimeric simian-human immunodeficiency virus (SHIV) challenge in rhesus macaques (*Macaca mulatta*). However, a major problem in establishing vaccine-induced nAb protection in the SHIV/macaque model has been the notorious difficulty in inducing nAbs by immunization. Indeed, induction of broadly neutralizing antibodies (bnAbs) via immunization has thus far only been achieved reproducibly in cows (Sok et al., 2017). However, we recently showed reliable induction of autologous strain-specific nAbs in macaques against a hard-to-neutralize Tier 2 HIV isolate through use of well-ordered and stabilized HIV envelope glycoprotein (Env) SOSIP trimers as immunogens in optimized approaches (Pauthner et al., 2017), building on previous SOSIP immunization studies in NHPs (Havenar-Daughton et al., 2016; Sanders et al., 2015; Torrents de la Peña et al., 2017). To carry out a protection experiment in macaques then requires construction of a SHIV with the same Env sequence as the immunizing trimer. Fortunately, it has recently become possible to reliably generate infectious SHIVs using *env* sequences from most primary Tier 2 HIV strains (Del Prete et al., 2017; Li et al., 2016).

In this study, building upon the advances in both trimer-based immunization strategies and SHIV generation, we immunized macaques with SOSIP trimers of the BG505 *env* sequence (de Taeye et al., 2015; Kulp et al., 2017; Torrents de la Peña et al., 2017), induced BG505-specific Tier 2 nAbs, and then challenged animals intrarectally with the neutralization-resistant, pathogenic SHIV_BG505_ (Li et al., 2016), that carries the S375Y mutation to increase infectivity in NHPs. We found that protection could be achieved and was critically dependent on the level of serum nAb titers, but not on other antibody parameters such as V3 binding titers, antibody-dependent cellular cytotoxicity (ADCC), or the induction of T cell activity. We determined an approximate threshold titer for vaccine-induced protection that establishes an experimental benchmark for comparison with nAb-based vaccines to HIV-1.

## RESULTS

### Balanced selection of challenge animals

Our goal was to assess the capability of vaccine-elicited Tier 2 nAbs to protect from homologous Tier 2 challenge with neutralization-resistant, pathogenic SHIV_BG505_ (Li et al., 2016). We previously developed a protocol for the reliable induction of nAbs and immunized 78 NHPs (Pauthner et al., 2017), inducing varying levels of autologous Tier 2 nAb titers after three immunizations with native-like BG505 Env trimers (de Taeye et al., 2015; Kulp et al., 2017; Torrents de la Peña et al., 2017). To design a challenge study powered to detect differences between NHPs with either high or low BG505 nAb titers, we selected six NHPs that were among the top neutralizers and carefully matched them as closely as possible, in terms of gender, age and weight, with six low nAb titer animals that received similar or identical immunogens (Figure S1, A to C). We note that none of the protective or viral breakthrough or antibody kinetic effects described below could be associated with a particular immunogen; as will be seen, observed effects are primarily associated with nAb titer. We further enrolled 12 unimmunized control animals into the study. All animals were genotyped for Mamu and TRIM-5α alleles associated with host restriction in non-human primates (Table S1).

### Design of the SHIV_BG505_ challenge study

To identify a challenge dose that reliably infects unimmunized control animals, we performed a pilot study by intrarectally (IR) inoculating two groups of six macaques at weekly intervals with either 0.5 x 10^8^ or 1.4 x 10^7^ virions of the SHIV_BG505_ S375Y challenge virus grown in rhesus CD4^+^ T cells (Figure S2). For the main study, we selected a challenge dose of 1.4 x 10^7^ virions (1ml of 1:75 diluted challenge stock), since it infected at least 4/6 animals following the 1^st^ challenge and the remaining two animals after the 2^nd^ challenge in the pilot study. To maximize nAb titer levels in NHPs prior to challenge, high and low nAb titer animals each received a fourth immunization with the previously used immunogens, adjuvanted with an ISCOMATRIX-like saponin (Figure 1A). All NHPs responded with increased autologous nAb titers two weeks post-boost. High and low nAb titer animals continued to show significantly different geometric mean ID_50_ titers of 1:3790 and 1:103 to BG505 S375Y pseudovirus (p=0.002, Figure 1B), respectively. Neutralization titers to rhesus CD4^+^ T-cell grown SHIV_BG505_ S375Y challenge stock were ~30-fold lower, with significantly different geometric mean titers of 1:102 and < 1:10 when tested on TZM-bl target cells (p=0.002, Figure 1C), respectively.

**Figure 1.**
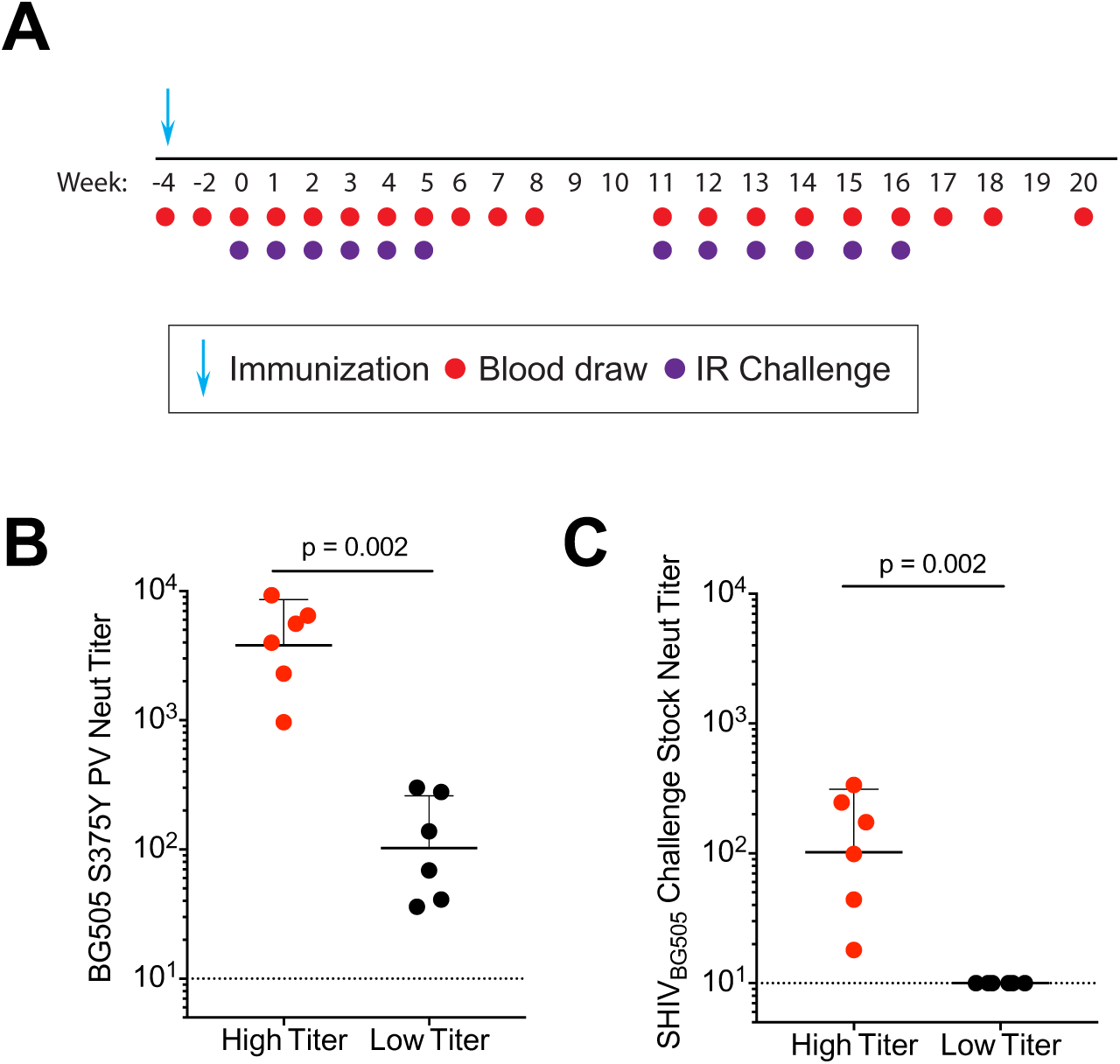
Challenge study design. (A) Animals, except for the controls, received a booster immunization using the same immunogen that had last been used during the preceding immunization study (Pauthner et al., 2017), typically 100µg SOSIP trimer adjuvanted in IscoMIT. Intrarectal (IR) challenges with SHIV_BG505_ S375Y commenced 4 weeks thereafter. All groups of animals received 6 IR challenges starting at week 0. High nAb titer animals that had undetectable serum viral loads at week 6 received a second set of 6 weekly IR challenges starting week 11. (B-C) Serum neutralizing ID_50_ titers in high and low nAb titer animals at week −2: BG505 S375Y pseudovirus (B) and rh-CD4-grown SHIV_BG505_ S375Y challenge stock. (C). Shown are geometric means with geometric SD, significant differences were determined using two-tailed Mann-Whitney U tests.

### Robust protection of high nAb titer group NHPs

Four weeks after the booster immunization, all animals received six weekly IR challenges with SHIVBG_505_. To maximize comparability, viral loads for all animals and time points were simultaneously measured at weeks 6 and 20 (Figure 2, A to C). 5/6 concurrent unimmunized control animals were infected after the 1^st^ challenge and the remaining animal became viremic after the 2^nd^ challenge (Figure 2A). Combined with the unimmunized control NHPs of the dose-matched titration group (Figure S2), at least 9 of 12 unimmunized animals became infected following a single challenge, which approximates to an animal infectious dose of 75% (AID75) (Table S2). Thus, the dose of 1.4 x 10^7^ SHIV_BG505_ virions per IR inoculation employed in this study sets a relatively high bar for protection. Unimmunized control animals showed high peak viremia (geometric mean of 5.5 x 10^6^ copies/ml) and consistent set point viral loads in the range of 9.8 x 10^2^ – 4.7 x 10^4^ (geometric mean of 6.2 x 10^3^) at 12 weeks post-infection (Figure 2, A and E, Figure S2).

**Figure 2.**
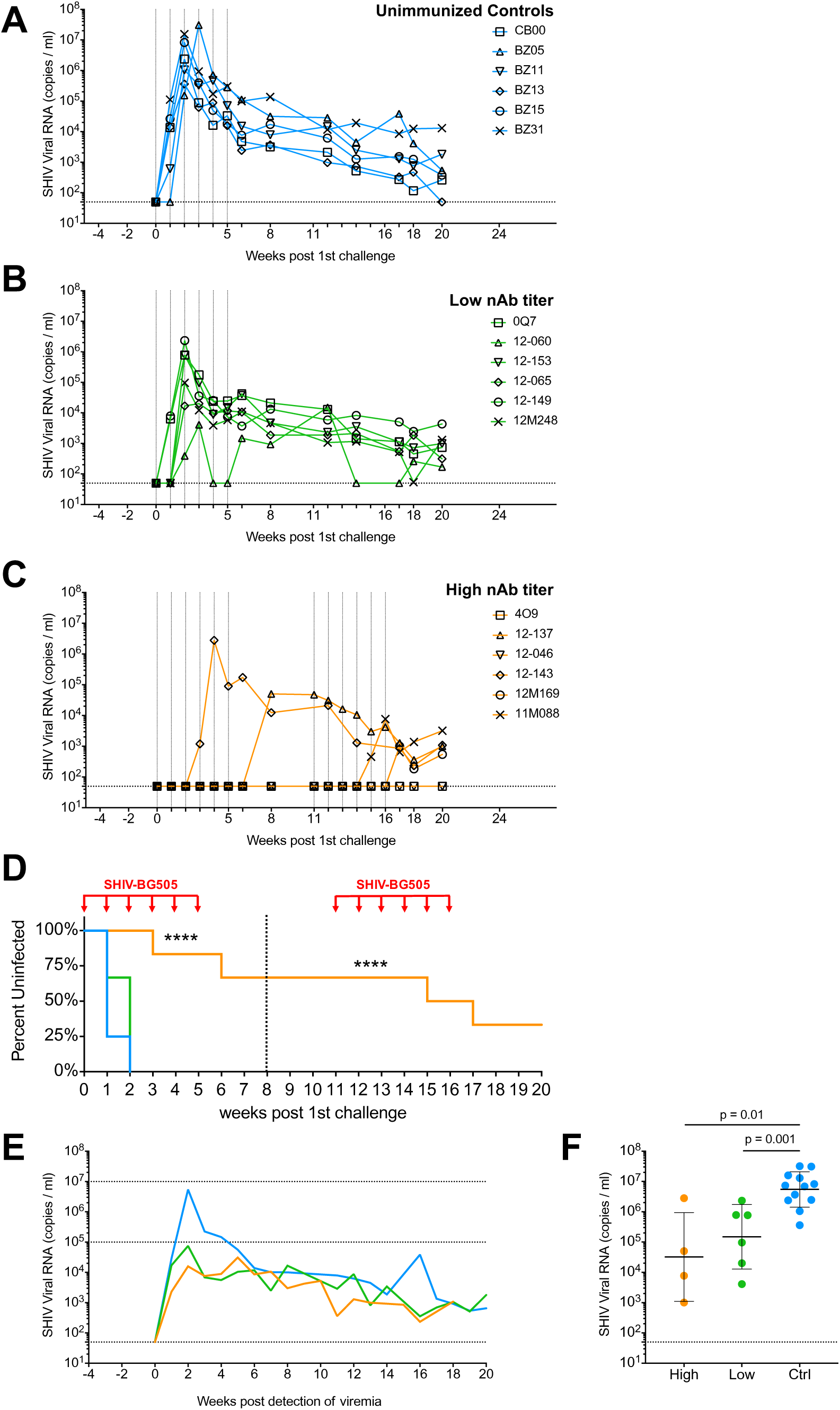
High nAb titer animals show robust protection. Viral loads of animals throughout the challenge schedule: unimmunized concurrent controls (**A**), low nAb titer (**B**) and high nAb titer macaques (**C**). IR challenges are indicated with vertical dotted lines. Horizontal dotted lines denote the limit of detection. (**D**) Kaplan-Meier curves indicating percent uninfected animals over the duration of the study. Challenge time points are indicated with red arrows. The high nAb titer group infection-rate was statistically different from the low nAb titer and unimmunized control groups. **** = p < 0.0001. Statistics were calculated for both the first (dotted line at week 8) and second challenge sets (see Table S3) (**E**) Geometric mean viral loads of indicated groups, normalized to the detection of viremia in the blood. Horizontal lines at 10^5^ and 10^7^ viral RNA copies/ml serve as visual aids. (**F**) Comparison of peak viral loads between high nAb titer (high), low nAb titer (low), and unimmunized (ctrl) animals. Geometric mean titers are shown with geometric standard deviations. Significant differences were determined using two-tailed Mann-Whitney U tests.

2/6 low nAb titer animals became infected after the first challenge and the remaining four animals became viremic following the second challenge (Figure 2B), indicating that low nAb titer animals had a possible mild reduction in per-exposure risk compared to unimmunized controls, but the difference was not significant (Figure 2D, Table S3). However, low nAb titer animals had significantly lowered peak viral loads compared to unimmunized controls (1.5 x 10^5^ vs. 5.5 x 10^6^ copies/ml) (p = 0.001, Figure 2, E and F).

In contrast, high nAb titer animals showed highly significant protection from challenge following the first set of challenges at week 8 (Figure 2D, Table S3). Except for macaque 12-143, no animals showed viremia at week 6 and were therefore scheduled to receive a second set of six challenges starting at week 11. The goal of the second challenge set was to assess the duration of protection and to estimate a protective nAb titer threshold as nAb titers declined over time. Over the course of both challenge sets, four initially high nAb titer animals became viremic, after 3, 6, 10 and 12 virus inoculations; however, two animals showed complete sterilizing protection (Figure 2C). In addition, infected high-titer macaques showed significantly lowered peak viremia compared to unimmunized controls (3.2 x 10^4^ vs 5.5 x 10^6^ copies/ml; p = 0.01, Figure 2, E and F), similar to the low nAb titer animals. We theorize that sub-protective levels of serum nAbs at the time of infection, as well as activation of vaccine-induced memory B cells leading to the rapid production of Abs, likely curtail emerging primary viremia, thus reducing peak viral loads.

The protection from infection for high nAb titer animals was highly significant compared to unimmunized controls after both 6 and 12 challenges (p < 0.0001, Figure 2D, Table S3) and animals in this group remained uninfected for a median of 11 challenges (Table S3). It should be emphasized that for all vaccinated animals, nAb titers declined throughout the challenge schedule, unless animals became infected as detailed below. In this respect, our study distinguishes itself from those in which antibody titers leveled off prior to challenge, as a result of the short 4-week interval here between final immunization and first challenge. However, we deliberately took advantage of declining nAb titers to determine a nAb-mediated threshold of protection.

### Tier 2 nAb titers correlate with protection

Unimmunized control animals developed BG505 S375Y pseudovirus ID_50_ nAb titers 8-12 weeks postinfection in response to SHIV_BG505_ S375Y infection (Figure 3A). By comparison, vaccine-induced nAb titers in low titer animals initially declined, but then began to rise only 1-2 weeks postinfection, i.e. much more rapidly than in unimmunized animals (Figure 3B). The early rise of nAb titers following infection of low nAb titer animals is thus likely due to recall responses of BG505 Env immunogen-induced memory B cells. Interestingly, BG505 nAb titers rose to substantially higher ID_50_ titers (3/6 animals >1:750) than previously achieved by four immunizations of these six animals with ISCOMs-adjuvanted BG505 native-like Env trimers (Figures 1B, 3B, S1B). The marked increases in BG505 nAb titers following infection suggest that outbred macaques that did not respond well to vaccination were not inherently incapable, by genetic or other means, of developing high nAb titer responses, although this conclusion should be caveated by the observation that antigen dose and delivery vary greatly between vaccination and natural infection. Better immunogen presentation and more targeted adjuvants are likely needed to increase the reliability of high nAb titer development, and to address current shortcomings in the durability of nAb responses induced by protein-only immunizations (Havenar-Daughton et al., 2017).

**Figure 3:**
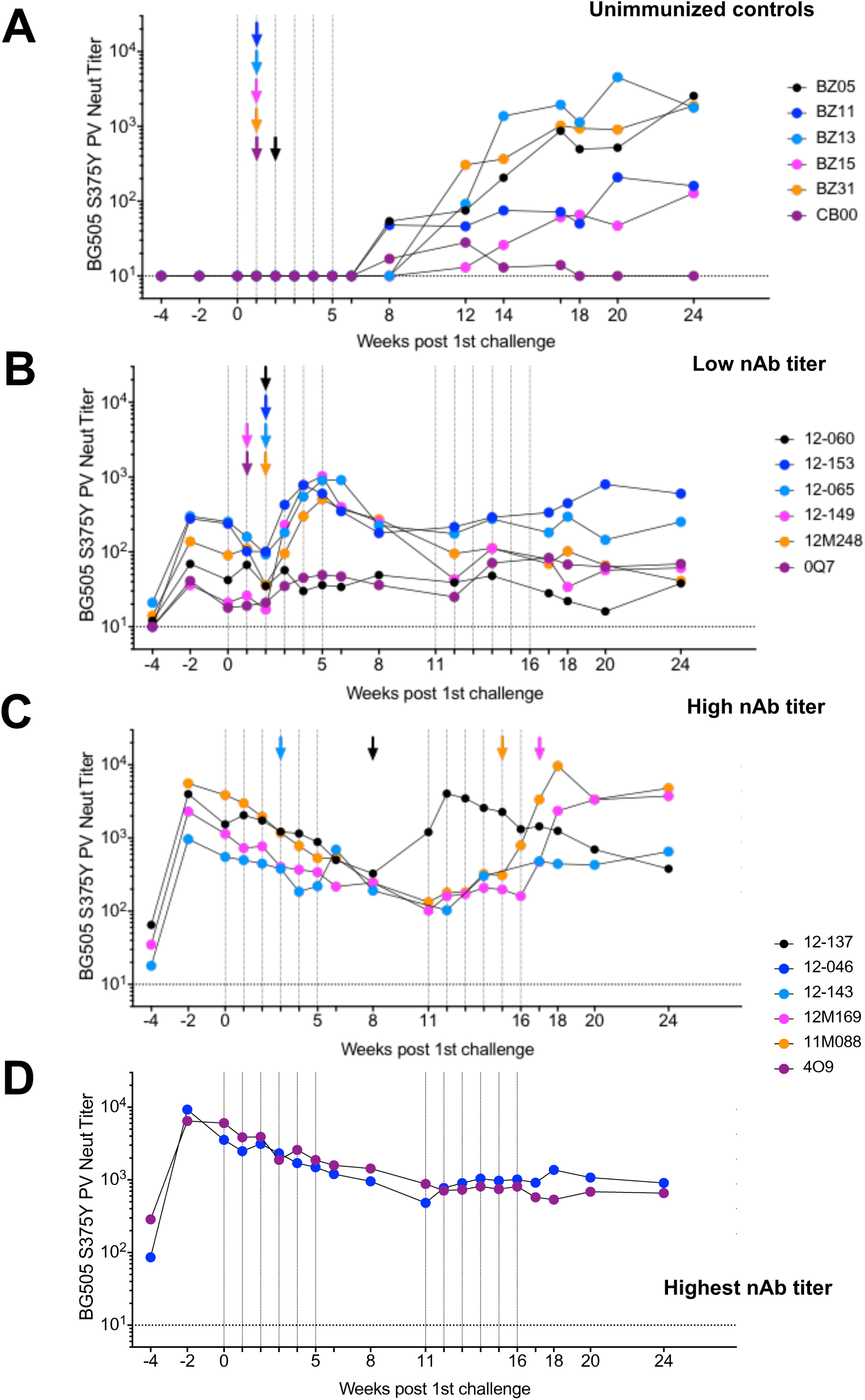
Longitudinal development of autologous Tier 2 nAb titers. (**A-D**) Serum neutralizing antibody titers throughout the challenge schedule: BG505 S375Y pseudovirus ID_50_ nAb titers rise 8-12 weeks post infection in unimmunized animals (**A)** or 1-2 weeks following detection of viremia in low nAb titer animals (**B**). BG505 S375Y pseudovirus ID_50_ nAb titers in macaques that became infected over time (**C**) or showed sterilizing protection (**D**). In (**D**), nAb titers peaked at week −2 following a final boost at week −4 and slowly declined until ~week 10 after which titers plateaued and remained stable for the duration of the study. Macaques infected during the second set of challenges displayed similar nAb titer plateaus as protected animals until infection. Animals that became infected showed a surge in nAb titer followed by a slow decline e.g. animal 12-137. First detection of plasma viremia is indicated by colored arrows corresponding to the animal IDs shown in the respective figure legends.

High nAb titer animals that became infected showed a comparable increase in BG505 S375Y nAb titers ~1-4 weeks following infection. The only exception was animal 12-137, who suppressed viremia for 3 weeks following challenge at week 5, and thus delayed a surge in nAb titers until week 11 (Figure 3C). Animal 12-143, which became viremic at week 3, showed only a small rise in nAb titers at week 6, suggesting possible rapid viral escape. PacBio sequencing of viral species in 12-143 plasma at week 8 in fact revealed that >95% of sequenced *env* genomes contained putative escape mutations at residues 168 and 192 (Figure S3A). Similarly, *env* genomes in 12-137 plasma at weeks 12 and 16 showed putative escape mutations at residues 354 and 356, flanking the N355 glycan, which coincided with onset of nAb titer decay at week 12 (Figure S3B, Figure 3C). NAb specificities to the N355-region were observed in BG505 SOSIP immunized rabbits (Klasse et al., 2018), and were detected in week 0 plasma of animal 12-137 using electron microscopy based serum mapping (Figure S3C) (Bianchi et al., 2018). We could further show that the observed viral point mutations do confer neutralization resistance to sera from the respective animals (Figure S3D-E). Animals 12M169 and 12M088, which become infected at weeks 16 and 14, respectively, exhibited slow declines in vaccine-induced nAb titers which then rose following infection (Figure 3C). The nAb titers of fully protected animals 12-046 and 4O9 (Figure 3D) initially declined and then plateaued at ~1:800 around week 10 and remained stable for the remainder of the study. This trend was mirrored in longitudinal ELISA EC_50_ binding titers. (Figure S3F-G). Uninfected animals retained robust nAb titer levels over 1 year past the final immunization (Figure S3H).

The differences in both BG505 S375Y pseudovirus as well as SHIV_BG505_ challenge stock neutralization ID_50_ titers between high and low nAb titer animals at week −2 were, as anticipated, highly significant (Figure 1, B and C). Peak nAb titers at week −2 accurately predicted the duration of protection, identifying nAb titers as the primary correlate of protection (p < 0.0001, Figure 4A). Using the BG505 S375Y pseudovirus assay, a statistically significant difference was found between nAb titers in immunized animals 7d prior to onset of viremia and animals that remained uninfected until week 20 (p = 0.03, Figure 4B, Figure S4A).

**Figure 4.**
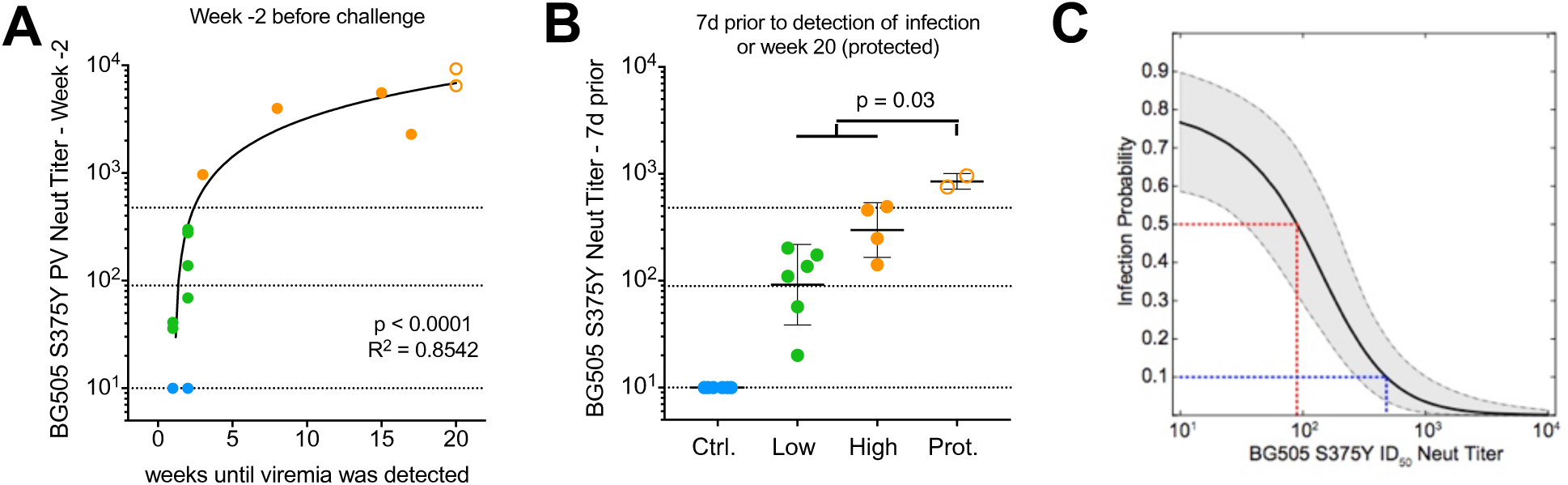
Protection is associated with serum nAb titers greater than ~1:500. (**A**) BG505 S375Y pseudovirus ID_50_ nAb titers at week −2 predict and correlate with the duration of protection. (**B**) BG505 S375Y pseudovirus ID_50_ nAb titers of control (Ctrl.), low nAb titer (Low) and high nAb titer (High) animals 7 days before detection of viral load in the blood and at week 20 for protected (Prot.) animals that showed sterilizing protection throughout the study. All nAb titers were measured in TZM-bl assays. Correlations were calculated using Spearman correlation tests, comparisons between groups were calculated using Mann-Whitney U tests. Horizontal lines indicate 50% and 90% protective nAb titers as defined in 3G. (**C**) The 5%, median, and 95% credible intervals (CI) are shown for the probability of infection in relation to serum BG505 S375Y pseudovirus nAb titer, inferred using a modified Bayesian logistic regression model (see figs. S10-S12). The posterior median infection probability at the limit of nAb titer detection was 77%, agreeing closely with an estimated animal infectious dose of 75% in unimmunized animals. A median infection probability of 50% is attained with an ID_50_ titer of 1:90 (red line, CI: 34-178), and an infection probability of 10% with an ID_50_ titer of 1:476 (blue line, CI: 272-991).

To numerically quantify the relationship between BG505 S375Y pseudovirus nAb titers and likelihood of infection, we developed a modified Bayesian logistic regression model using the neutralization and viral load data from all three animal groups (Figure 4C, S5). The posterior median infection probability at the limit of nAb titer detection was 77%, agreeing closely with an estimated animal infectious dose of 75% in unimmunized controls. A median per-challenge infection probability of 50% was attained with ID_50_ titers of 1:90, which agrees well with the often-quoted 50% protective ID_50_ titer of ~1:100, derived from bnAb passive transfer studies (Hessell et al., 2018; Moldt et al., 2012; Parren et al., 2001; Pegu et al., 2014; Shingai et al., 2014), although we note that various different neutralization assays with differing sensitivities were employed in these studies. To achieve an infection probability of 10%, or 90% protection, an ID_50_ titer of 1:476 (CI: 272-991) was required. In agreement with our model, animals with nAb titers above ~1:500 remained protected over all 12 challenges, while animals with nAb titers below 1:200 generally became infected with only 1-2 challenges. For the rhesus CD4^+^ T cell grown SHIV_BG505_ S375Y virus stock, an ID_50_ titer of ~1:30 (Figure S4B) was the protection threshold. The observed disparity between pseudovirus and PBMC-grown virus assays was relatively large compared with that reported for many mAbs but still within previously observed ranges (Figure S4C) (Cohen et al., 2018; Provine et al., 2012). Thus, Tier 2 nAb titers both predicted and correlated with protection from infection.

### T cell activity and serum antibody-dependent cell-mediated cytotoxicity (ADCC) do not correlate with protection

We further investigated other parameters that may have contributed to protection. Robust Env-specific CD4^+^ T cell responses were elicited by BG505 Env trimer immunization and were equivalent in magnitude between the high and low nAb titer groups of immunized animals before challenge (Figure 5, A and E, Figure S6, A to C). Cytokine-producing Env-specific CD4^+^ T cells were also comparable between the two groups of immunized animals before challenge (IFNgγ^+^, Figure 5, B and F; TNF^+^CD40L^+^, Figure 5, C and G, Figure S6D) and protein vaccine-elicited Env-specific CD8^+^ T cells were undetectable, as expected (Figure 5,D and H). Thus, Env-specific CD4^+^ T cells or CD8^+^ T cells are not a correlate of protection.

**Figure 5.**
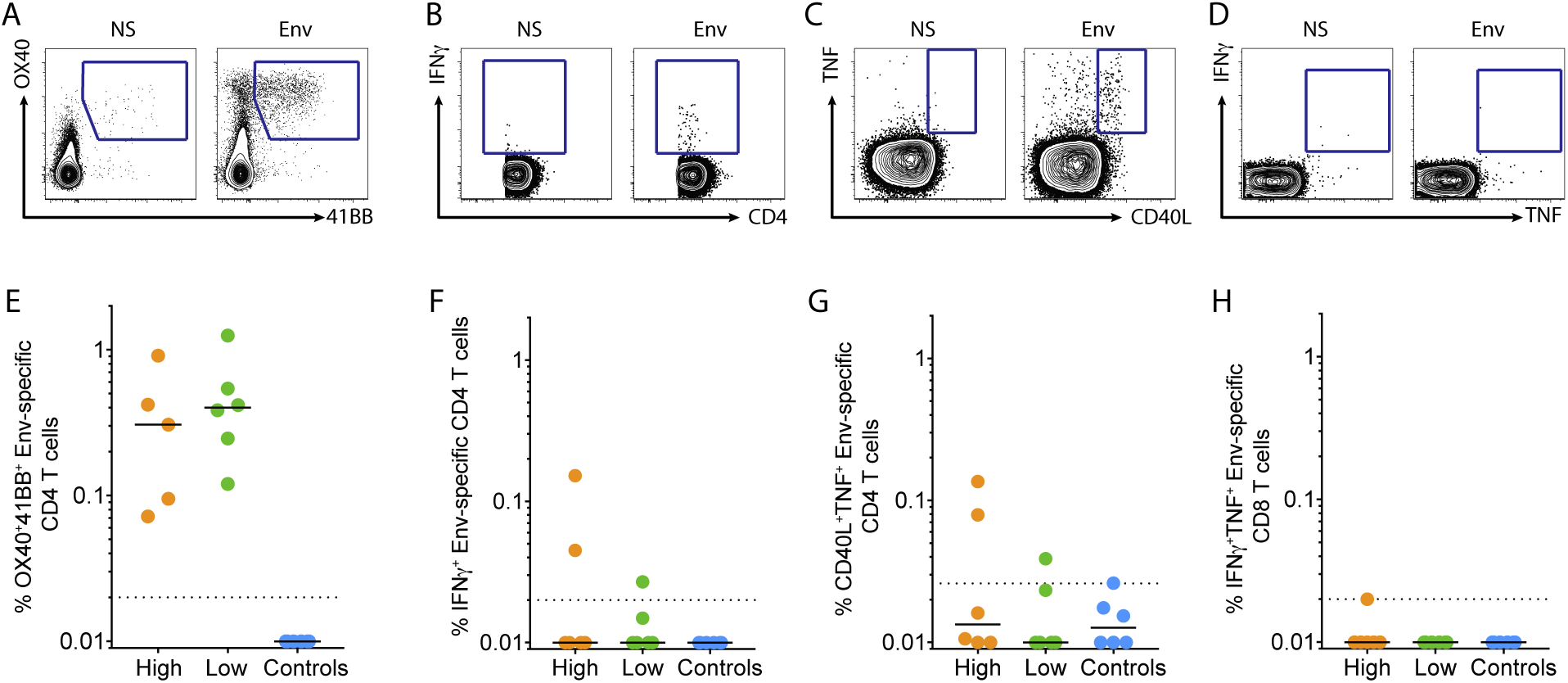
HIV Env-specific CD4^+^ T cells and Env-specific CD8^+^ T cells at week 0 are not associated with the observed protection from infection. (**A-C**) Representative flow plots of Env-specific CD4 T cells from week 0 PBMCs: using an OX40/4-1BB AIM assay (Reiss et al., 2017) (**A**), ICS assay for IFNγ (**B**), and ICS assay for TNF/CD40L (**C**) when not stimulated (ns) versus stimulated with antigen (Env). (**D**) Representative flow plot of IFNγ and TNF expression in CD8 T cells by ICS when not stimulated (ns) versus stimulated with antigen (Env). (**E-G**) Quantification of the percent of CD4 T cells that are Env-specific based on: OX40/4-1BB (**E**), IFNγ (**F**), or CD40L/TNF (**G**) expression. (**H**) Quantification of the percent of CD8 T cells that are Env-specific based on IFNγ and TNF expression. Signal from the unstimulated condition was subtracted from the antigen-specific signal for each sample. Each dot represents an individual animal.

Concerns have been raised about vaccine-elicited CD4^+^ T cell responses enhancing susceptibility to infection by HIV (Fauci et al., 2014; Hu et al., 2014) or SIV (Fouts et al., 2015; Staprans et al., 2004) by providing more targets for infection at the mucosal site of transmission (Bukh et al., 2014; Carnathan et al., 2015; Martins and Watkins, 2017; Qureshi et al., 2012), most likely due to the presence of activated Th1 cells in the mucosa, which was correlated with CCR5, α4β7, or proliferation in different studies. Minimal Th1 cells were detected in the BG505 Env trimer immunized animals (IFNg^+^ CD4^+^ T cells, Figure 5, B & F). CCR5^+^, Ki67^+^ or Ki67^+^/α4β7^+^ CD4^+^ T cells in peripheral blood prior to challenge were not correlated with susceptibility to infection or protection (Figure S6, E to H). Thus, we observed robust protection of high nAb titer animals against a mucosal SHIV challenge despite substantial levels of Env-specific vaccine-induced CD4^+^ T cells in peripheral blood at 4 weeks after the final immunization. The difference in our study may be due to a lack of Th1 or mucosal homing CD4+ T cells in response to the protein vaccine, compared to previously used viral vectors (Bukh et al., 2014; Carnathan et al., 2015; Fauci et al., 2014; Hu et al., 2014; Staprans et al., 2004). Alternatively, nAb-mediated protection against HIV/SIV may more readily overcome possible adverse consequences of increased numbers of activated CD4^+^ T cell targets than the non-neutralizing Abs (nnAbs) raised in the earlier studies.

To investigate possible contributions of ADCC of both nAbs and nnAbs, we tested animal sera in two infection-based assays; SHIV_BG505_-infected CEM.NKR luciferase reporter cells (Alpert et al., 2012) (Figure 6A) and flow cytometric analysis of ADCC in p27^+^ SHIV_BG505_-infected CEM.NKR target cells (Veillette et al., 2014) (Figure 6B). Using either assay, we failed to detect meaningful ADCC activity at week 0 with the exception of a single animal, 12-149, which was a low titer animal whose ADCC activity was non-specific and included activity against control SIV_mac_239 (Figure S7A). The absence of observed ADCC activity can be partially explained by the Tier 2 character of BG505 Env. In native Env trimer-based ADCC assays, nnAb and Tier 1 nAbs fail to mediate ADCC-activity against, hard-to-neutralize Tier 2 HIV isolates, as previously reported (Bredow et al., 2016; Ding et al., 2016). In addition, ADCC activation in infection-based assays varies strongly depending on the targeted epitope, which is likely related to the Ab binding stoichiometry to the epitope and the ability to cross-link sparse trimers on the virion surface (Figure S7B-D) (Bredow et al., 2016; Ding et al., 2016).

**Figure 6:**
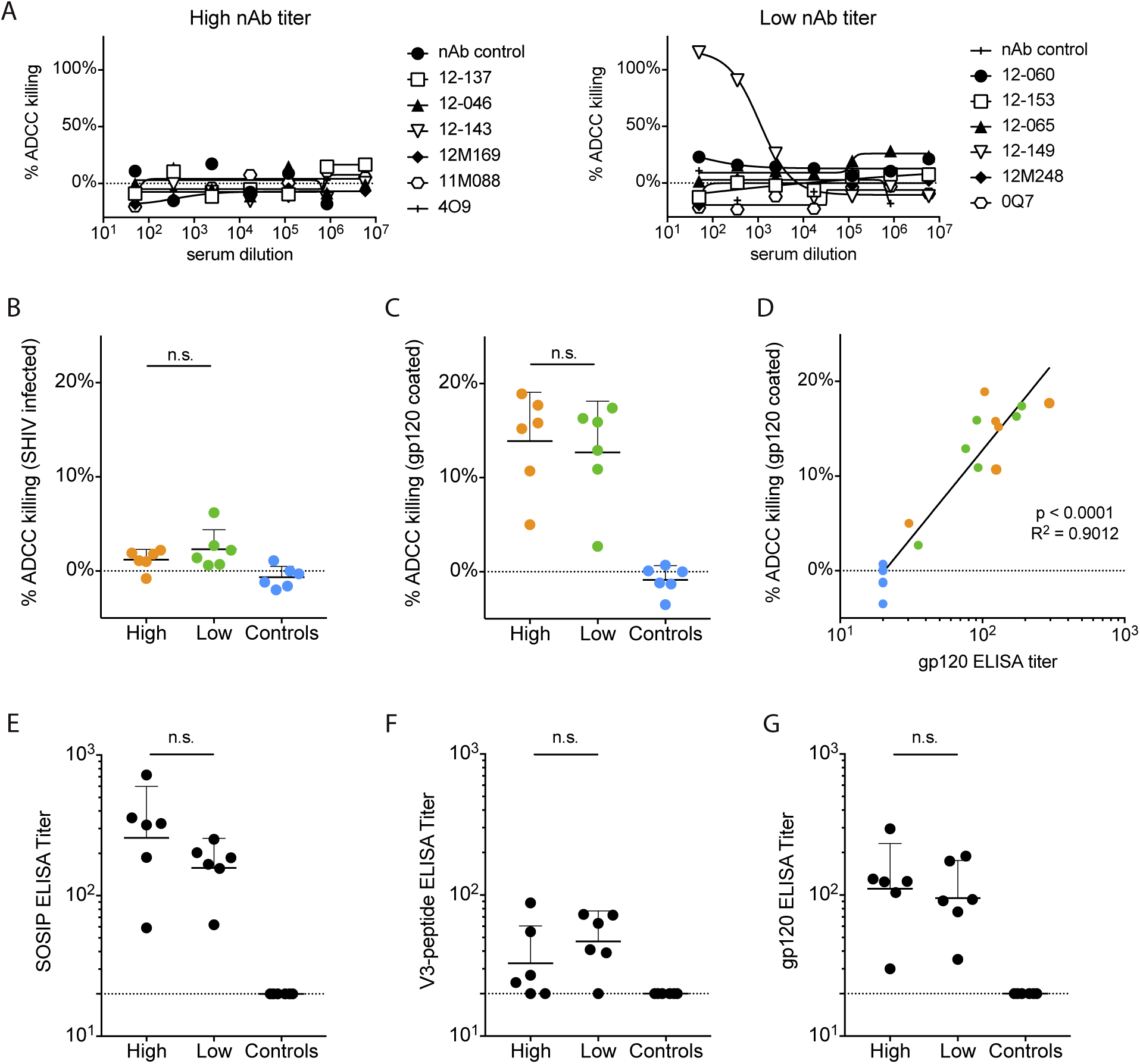
ADCC activity at week 0 measured in SHIV-infection as well as gp120-based assays is not associated with the observed protection from infection. (**A-G**) ADCC-activity from sera of high and low nAb titer as well as control animals at week 0. ADCC activity in titrated sera was measured using SHIV_BG505_ challenge stock infected CEM.NKR luciferase-reporter target cells and CD16 transfected KHYG-1 effector cells (**A**) or in 1:250 diluted sera by flow cytometric analysis of ADCC activity in either p27^+^ SHIV_BG505_-infected CEM.NKR cells (**B**) or BG505 g120-coated CEM.NKR cells (**C**), using PBMCs as effector cells. ADCC-activity in BG505 gp120-coated CEM.NKR cells correlated with BG505 gp120 binding titers (**D**). (**E-G**) ELISA EC_50_ binding titers to: BG505 SOSIP.664 (**E**), BG505 V3-peptide (**F**) or BG505 gp120 (**G**). Sera from high and low titer animals, as well as unimmunized control animals were tested for ELISA binding titers at week 0. Correlations were calculated using Spearman correlation tests, comparisons between groups were calculated using Mann-Whitney U tests.

Unlike infection-based assays, ADCC killing measured on CEM.NKR target cells coated with BG505 gp120 was robust, but did not distinguish between high and low nAb titer animals and, therefore, was not associated with protection (Figure 6C). ADCC killing of gp120-coated cells did correlate with BG505 gp120 binding, indicating that gp120-binding antibodies are sufficient to induce ADCC in this assay (Figure 6D), but cannot mediate ADCC to native membrane-bound Env on infected cells. Thus, ADCC unlikely contributes to protection. We also observed considerable staining of p27^-^ uninfected bystander T cells by both mAbs and animal sera, which appears to result from antibody binding to shed gp120 from infected cells that is captured on CD4 of uninfected cells (Figure S7D) (Richard et al., 2018). Overall, these results suggest caution in the use of ADCC assays that are either based on recombinant gp120 or gp140 binding, rather than native Env on virus-infected cells, or cannot distinguish productively infected from uninfected bystander cells (Ackerman et al., 2016; Ferrari et al., 2011; Huang et al., 2016; Johansson et al., 2011; Kristensen et al., 2018). We note that the results pertain to ADCC; there remains the possibility that other Fc-mediated effector functions might contribute to protection.

Lastly, we determined BG505 SOSIP.664 (Figure 6E), V3-peptide (Figure 6F), and BG505 gp120 binding titers (Figure 6G) for all groups at week 0 since V3-targeting antibodies (Balasubramanian et al., 2018), and binding antibodies in general have been associated with anti-viral activities (Excler et al., 2014). No significant differences between high and low neutralizer animals were detected.

## DISCUSSION

Vaccine protection against HIV in humans and against SIV and SHIV in macaques has been associated with non-neutralizing antibodies (Barouch et al., 2015; Haynes et al., 2012). Here, we demonstrate that vaccine-induced Tier 2 nAbs, but not other antibody parameters such as V3 binding titers, antibody-dependent cellular cytotoxicity (ADCC), or induction of T cell activity, are a correlate of protection from homologous SHIV_BG505_ infection in macaques. Notably, we employed a challenge dose of virus corresponding to an AID75, which sets a relatively high bar for protection, given that most animals (~53%) in the control arm were estimated to have been productively infected by two or more viruses (Table S2). Similar rates of multivariant virus transmission have been reported in men who have sex with men and injection drug users who acquire HIV-1 infection (38% and 60% with a MOI of 2 or higher, respectively), while heterosexual cohorts show lower multivariant transmission frequencies (~19%). Thus, our model mimics the conditions of productive transmission events, underlining the physiological relevance of the challenge dose that we used (Bar et al., 2010; Li et al., 2010).

We show, in the model system described, that animals remain protected from SHIV infection in a nAb titer-dependent manner, which suggests a strong relationship between circulating nAb titers in the blood and protection from mucosal challenge with difficult-to-neutralize, Tier 2 SHIV_BG505_. At the same time, our data suggest that vaccine protection can occur in the absence of ADCC. We show that unprotected animals have relatively high levels of ADCC when measured in a widely used ADCC assay that uses target cells coated with monomeric gp120, but not with SHIV_BG505_ infected target cells. We further provide evidence that adjuvanted protein immunization with HIV Env can induce nAb titers that are durable and protective over longer periods of time, if high initial nAb titers following immunization can be reached. This has been a major concern in the HIV vaccine field (Sundling et al., 2013), but also for other protein-based vaccines, such as recombinant influenza vaccines (Krammer and Palese, 2015). Importantly, we identified that a serum ID_50_ nAb titer of ~1:500 against the homologous BG505 S375Y pseudovirus at the time of challenge can confer reliable protection of > 90%, meaning that 9 of 10 challenges with a physiologically-relevant AID75 dose would not result in infection. Finally, protection is observed for polyclonal neutralizing Ab responses that, as above and earlier {Pauthner:2017fd}, target multiple specificities on Env and not simply the previously described glycan hole on BG505 Env.

In conclusion, we provide evidence that protein immunization with native-like Env trimers can induce potent and protective nAb titers in the SHIV/macaque model. Thus, nAb-mediated protection from Tier 2 virus challenge is not limited to bnAbs, which are generally focused to a single site of vulnerability and have a defined effector-function profile, but can also be accomplished by polyclonal autologous nAb responses of sufficient magnitude and specificity, which comprise a broad range of neutralizing and non-neutralizing antibody lineages to various, often overlapping epitopes that interact in both synergistic or competitive ways (Klasse et al., 2018; Pauthner et al., 2017; Sanders et al., 2015; Torrents de la Peña et al., 2018). The protective nAb titer threshold against the homologous challenge virus that we determined is in rough accord with passive antibody transfer studies and provides a benchmark for comparison with upcoming antibody protection studies against HIV in humans (www.ampstudy.org).

## ACKNOWLEDGEMENTS

We would like to thank Laura Pruyn and Joan Allmaras for excellent administrative support. The funding for this study was provided by NIAID UM1AI100663 (Center for HIV/AIDS Vaccine Immunology and Immunogen Discovery). The Bill & Melinda Gates (OPP1084519) and the International AIDS Vaccine Initiative helped support the design of some immunogens used in this study. This work was further supported by the National Institutes of Health grants AI121135 (DTE), AI124377, AI126603, AI128751 (DHB), OD011106 (Wisconsin National Primate Research Center) and grant OPP1145046 from the Bill & Melinda Gates Foundation (GMS). AF work was supported by CIHR foundation grant #352417. AF is the recipient of a Canada Research Chair on Retroviral Entry RCHS0235. JP is the recipient of a CIHR Fellowship Award. BM was supported by grants R00AI120851 and UM1AI068618 from the National Institute of Allergy and Infectious Diseases. PacBio SMRT sequencing was performed with the support of the Translational Virology Core at the UC San Diego Center for AIDS Research (P30 AI036214), and the IGM Genomics Center, University of California, San Diego, La Jolla, California.

## AUTHOR CONTRIBUTIONS

The TSRI CHAVI-ID immunogen working group consisting of SC, WRS, ABW, IAW, RTW and DRB, with the assistance of MGP, STB, GMS and DHB designed the challenge study and laid out the experimental strategy. JPN and DHB oversaw all rhesus macaque immunizations and challenges, including sample acquisition, processing, storage and distribution. PA and DHB performed and oversaw viral load assays. HL and GMS designed and produced the SHIV_BG505_ challenge stock. CAC, DWK and TT with oversight from DJI, ABW, WRS and RWS designed and produced the boosting immunogens for the study. MGP, RB, JHL and DRB designed HIV pseudovirus mutants and performed and oversaw neutralization experiments as well as ELISA binding experiments. CHD, SMR and SC performed and oversaw flow cytometric analysis of T cell activation. JP, RN and BVB with oversight from DTE, LH and AF performed ADCC assays. BN and MB with oversight from LH and ABW performed serum negative-stain EM analysis. BM and MGP performed statistical analysis of data sets. MGP, JPN, CHD, BM, SMR, JP, RN, BVB, TT, STB, DTE, LH, AF, IAW, RTW, DJI, WRS, ABW, RWS, SC, GMS, DHB and DRB analyzed data sets and contributed edits to the manuscript. MGP, SC and DRB wrote the manuscript.

## DECLARATION OF INTERESTS

The authors declare no competing interests

## STAR METHODS

## CONTACT FOR REAGENT AND RESOURCE SHARING

Further information and requests for resources and reagents should be directed to and will be fulfilled by Dennis Burton.

## EXPERIMENTAL MODEL AND SUBJECT DETAILS

### Rhesus macaques

Outbred Indian rhesus macaques (*Macaca mulatta*) were sourced and housed at Alphagenesis Inc, Yemasee, SC and maintained in accordance with NIH guidelines. These studies were approved by the appropriate Institutional Animal Care and Use Committees (IACUC). None of the NHPs were previously enrolled in other studies that are not explicity stated in the manuscript. All animals were genotyped for class I alleles Mamu-A*01, Mamu-B*08 and Mamu-B*17 and Trim5, which are associated with spontaneous virological control. Genotype and gender information for all animals is reported in Table S1. Additional information on high- and low-nAb titer group animals is published in Pauthner et al., 2017.

## METHOD DETAILS

### Rhesus monkey immunizations and challenge

Animals were immunized at 4 weeks before challenge (week −4) with a fourth dose of the previously administered immunogen for a given animal with adjuvant (Figure S1B) (Pauthner et al., 2017). The adjuvant used for this boost was an ISCOMATRIX-like nanoparticle comprised of self-assembled cholesterol, phospholipid, and Quillaja saponin prepared as previously described (Lovgren-Bengtsson et al., n.d.). All immunizations were administered as split doses. Each immunization consisted of two subcutaneous injections of 50 µg of Env trimer protein + 187.5 units (U) of saponin adjuvant, in sterile phosphate-buffered saline (PBS) diluent for a total of 100 µg of Env trimer protein + 375 U of Iscomatrix per immunization per animal. Subcutaneous immunizations were given in a volume of 0.5 ml with a 1 inch, 25-gauge needle at the medial inner mid-thigh of each leg. The subcutaneous injection technique consists of making a ‘skin tent’ and inserting the needle into the subcutaneous space at a 45° angle.

Serum was collected in SST Vaccutainer tubes (BD Biosciences) and processed according to the manufacturer’s instructions. Multiple aliquots of 0.5 ml were frozen at −80° C. Whole blood was collected in K2 EDTA Vaccutainer tubes (BD Biosciences) for plasma and PBMC isolation. Multiple aliquots of 0.5 ml of plasma were frozen at −80° C. PBMCs were isolated using Thermo Scientific Nunc EZFlip Conical Centrifuge Tubes per manufacturer’s instructions. PBMCs were isolated, counted, and re-suspended at 1 x 10^7^ cells/mL in FBS containing 10% DMSO. Aliquots were subsequently frozen in 1 mL vials using a Mr. Frosty freezing container (Nalgene, cooling rate of 1°C / minute) and placed in a −80° C freezer. The following day PBMC samples were moved to storage in a liquid nitrogen freezer tank.

Animals were atraumatically inoculated intrarectally with a 1:75 dilution of rhCD4-grown SHIV_BG505_ N332 S375Y ΔCT challenge stock (Li et al., 2016) in RPMI 1640 (Gibco), which amounted to 1.4 *10^7^ virions or 2 ng p27. See dataset S1B in Li et al. (Li et al., 2016) for a complete characterization of the challenge stock with respect to virion content and virion infectivity.

### Viral Load Assay

Plasma SHIV RNA levels in serum following infection were measured using a *gag-*targeted quantitative real-time RT-PCR assay as previously described (Hansen et al., 2013).

### Serum neutralization assays

Replication incompetent HIV pseudovirus was produced by co-transfecting *env* plasmids with an *env*-deficient backbone plasmid (pSG3Δ*env*) in HEK293T cells in a 1:2 ratio, using the X-tremeGENE 9 transfection reagent (Roche). Pseudovirus was harvested after 48-72 h by sterile-filtration (0.22 µm) of cell culture supernatants, and neutralization was tested by incubating pseudovirus and serum or mAbs for 1 h at 37 °C before transferring them onto TZM-bl cells as previously described (Pauthner et al., 2017). For replication competent SHIV_BG505_ neutralization, rhCD4-grown SHIV_BG505_ N332 S375Y challenge stock was used instead in a BSL3 facility with no further modifications.

Neutralization is measured in duplicate wells within each experiment. BG505 nAb titers for group comparisons were measured in three or more independent experiments that were subsequently averaged. The BG505 pseudovirus time course neutralization data shown in Figure 3 were generated in single large experiments, to test sera from all time points side-by-side, thus ensuring the highest nAb titer comparability between time points. Neutralization was tested starting at 1:10 serum dilutions followed by nine serial 3-fold dilutions to ensure the highest sensitivity and range of detection. Neutralization ID_50_ titers were calculated using the ‘One site – Fit logIC_50_’ regression in Graphpad Prism v7.0. ID_50_ nAb titers of incomplete neutralization curves that reached at least 50%, but less than 90% maximal neutralization, were calculated by constraining the regression fit through 0% and 100% neutralization, to ensure accurate calculation of half-way (50%) nAb titers. All neutralization titers are reported as ID_50_ titers. All nAb titer data panels show geometric mean titers with geometric SD. BG505 pseudovirus neutralization was tested using the BG505.W6M.ENV.C2 isolate (AIDS Reagents Program), carrying the T332N mutation to restore the N332 glycosylation site, as well as other indicated mutations that were added by site-directed mutagenesis.

### Serum binding ELISAs

Microlon 96-well plates (Corning) were coated overnight with streptavidin at 2.5 μg/mL (Thermo Scientific). Plates were then washed 4-5 times with PBS-tween (0.05%) and blocked with PBS + 3% BSA for 1 h at room temperature. If capturing biotinylated BG505 SOSIP.664-Avi or BG505-Avi gp120, proteins were added at 2.5 μg/mL in PBS + 1% BSA for 2 h at room temperature. For V3-peptide binding assays, no streptavidin was coated and instead BG505 V3-peptide (TRPNNNTRKSIRIGPGQAFYATG) was directly coated to Microlon 96-well plates at 2.5 μg/mL in PBS overnight. Plates were then washed 4-5 times with PBS-tween (0.05%) and serially diluted sera in PBS + 1% BSA were then added for 1 h at room temperature. Plates were then washed 4-5 times with PBS-tween (0.05%) and alkaline phosphatase-conjugated goat anti-human IgG (Jackson ImmunoResearch) was added for 1 h at a 1:1000 dilution (final concentration 0.33 μg/mL) in PBS + 1% BSA at room temperature. Plates were then washed 4-5 times with PBStween (0.05%) and absorption at 405 nm was measured following addition of phosphatase substrate in alkaline phosphatase buffer. We calculated half maximal EC_50_ binding titers using Graphpad Prism v7.0. All ELISA Ab data panels show geometric mean titers with geometric SD.

### ADCC assays

#### Luciferase-based CEM.NKR SHIV, HIV, SIV infection assay

ADCC activity was measured as previously described (Alpert et al., 2012). CEM.NKR-CCR5-sLTR-Luc cells, which express luciferase (Luc) upon infection, were infected with either HIV-1 BG505, SHIV BG505 or SIV_mac_239 by spinoculation in the presence of 40 µg/ml of polybrene. For HIV-1 BG505 and SHIV_BG505_ infections, *vif*-deleted infectious molecular clones were pseudotyped with Vesicular stomatitis virus G (VSVG). Two days post-infection with VSVG-pseudotyped HIV-1/SHIV_BG505_ and 4 days post-infection with SIVmac239, CEM.NKR-_CCR5_-sLTR-Luc cells were incubated at a 10:1 effector:target cell ratio either with an NK cell line expressing rhesus macaque CD16 in the presence of serial dilutions of rhesus macaque sera or an NK cell line expressing human CD16 in the presence of human monoclonal bnAbs. After an 8-hour incubation, Luc activity was measured using BriteLite luciferase substrate (PerkinElmer). Uninfected or infected cells incubated with NK cells in the absence of antibody or plasma were used to determine background and maximal Luc activity, respectively. The dose-dependent loss of Luc activity represents the antibody-dependent killing of productively infected target cells.

#### FACS-based CEM.NKR SHIV infection assay

VSVG-pseudotyped SHIV_BG505_ N332 S375Y virus was produced and titrated as previously described (Veillette et al., 2015). Viruses were then used to infect CEM.NKR-CCR5-sLTR-Luc cells by spin infection at 800 × *g* for 1 h in 96-well plates at 25 °C. Measurement of ADCC using the FACS-based assay was performed at 48h post-infection as previously described (Veillette et al., 2015). Briefly, infected CEM.NKR-CCR5-sLTR-Luc cells were stained with viability (AquaVivid; Thermo Fisher Scientific) and cellular (cell proliferation dye eFluor670; eBioscience) markers and used as target cells. Human PBMCs isolated from three different healthy HIV-uninfected individuals were used as effector cells and were stained with another cellular marker (cell proliferation dye eFluor450; eBioscience). Effector cells were added at an effector:target cell ratio of 10:1 in 96-well V-bottom plates (Corning, Corning, NY). A 1:250 final dilution of sera or 5 µg/ml of mAbs were added to appropriate wells and cells were incubated for 15 min at room temperature. The plates were subsequently centrifuged for 1 min at 300 g, and incubated at 37°C, 5% CO_2_ for 5 to 6 h before being fixed with a PBS-formaldehyde solution (2% formaldehyde final concentration). Cells were then permeabilized using the Cytofix/Cytoperm Fixation/Permeabilization Kit (BD Biosciences) and SHIV-infected cells were identified by intracellular staining using Alexa fluor 488-conjugated anti-p27 Abs (clone 2F12). Samples were analyzed on an LSRII cytometer (BD Biosciences). Data analysis was performed using FlowJo vX.0.7 (Tree Star). The percentage of ADCC was calculated with the following formula: (% of p27+ cells in Targets plus Effectors) − (% of p27+ cells in Targets plus Effectors plus Abs or sera) / (% of p27+ cells in Targets) by gating on infected living target cells. Of note, samples were deidentified and tested and analyzed blindly.

#### FACS-based gp120-coated CEM.NKR ADCC assay

CEM.NKR-_CCR5_-sLTR-Luc cells were coated with 1µg of recombinant HIV-1_BG505_ N332 gp120/million cells for 30 min at 37°C. gp120-coated target cells were used as target cells and were stained with viability (AquaVivid; Thermo Fisher Scientific) and cellular (cell proliferation dye eFluor670; eBioscience) markers. ADCC was performed as described above with the difference that after target/effector cells co-incubation, cells were fixed with a PBS-formaldehyde solution (2% formaldehyde final concentration) containing a constant number of flow cytometry particles (5×10^4^/ml) (AccuCount Blank Particles, 5.3 µm; Spherotech, Lake Forest, IL, USA). These particles are designed to be used as reference particles since their concentration is known, thus allowing to count the absolute cell number by flow cytometry. A constant number of particles (1×10^3^) were counted during cytometry acquisition in order to normalize the number of viable targets cells. Each sample was acquired with a LSRII (BD Bioscience, Mississauga, ON, Canada) and data analysis was performed using FlowJo vX.0.6 (Tree Star, Ashland, OR, USA). The percentage of ADCC was calculated with the following formula: (relative count of gp120-coated cells in targets plus effectors) - (relative count of gp120-coated cells in targets plus effectors plus Abs or sera) / (relative count of gp120-coated cells in targets) by gating live target cells (Veillette et al., 2015). Of note, samples were deidentified and tested and analyzed blindly.

### T cell analysis

Frozen aliquots of macaque PBMCs were thawed, washed once with RPMI + 10% FBS (R10), incubated with DNase (100ug/ml, StemCell Technologies 07900) for 15 minutes at 37C, then washed again and split in half for a CD8^+^ ICS assay and a CD4^+^ T cell Activation Induced Marker (AIM) assay (Reiss et al., 2017).

For the CD8+ T cell ICS assay, the sample was further split into three groups and either left unstimulated (NS), stimulated with BG505 Env peptides (5ug/ml), or stimulated with SEB (1ug/ml) for 2 hours at 37C. Brefeldin A was then added (2ug/ml), and the stimulations incubated for another 4 hours at 37C. The cells were then stained for 30 minutes at 4C with the fluorescent antibodies in the Surface Marker Panel below and washed twice with FACS buffer. They were fixed with eBio intranuclear fix/perm kit for 20 minutes, washed once with perm buffer, then stained with the antibodies in the Intranuclear Panel in perm buffer for 30 minutes at 4C. The samples were then washed with FACS buffer and acquired on a BD LSR Fortessa.

For the CD4^+^ T cell AIM assay, the sample was further split into three groups and either left unstimulated (NS) or stimulated with BG505 Env peptides (5ug/ml), or stimulated with SEB (100 pg/ml) for 24 hours at 37C. The cells were then stained for 60 minutes at 4C with the fluorescent antibodies in the AIM Surface Marker Panel below, washed with FACS buffer, fixed with 1% formaldehyde for 10 minutes at 4C, then washed again before acquisition on a BD LSR Fortessa.

CD8 Surface Marker Panel:

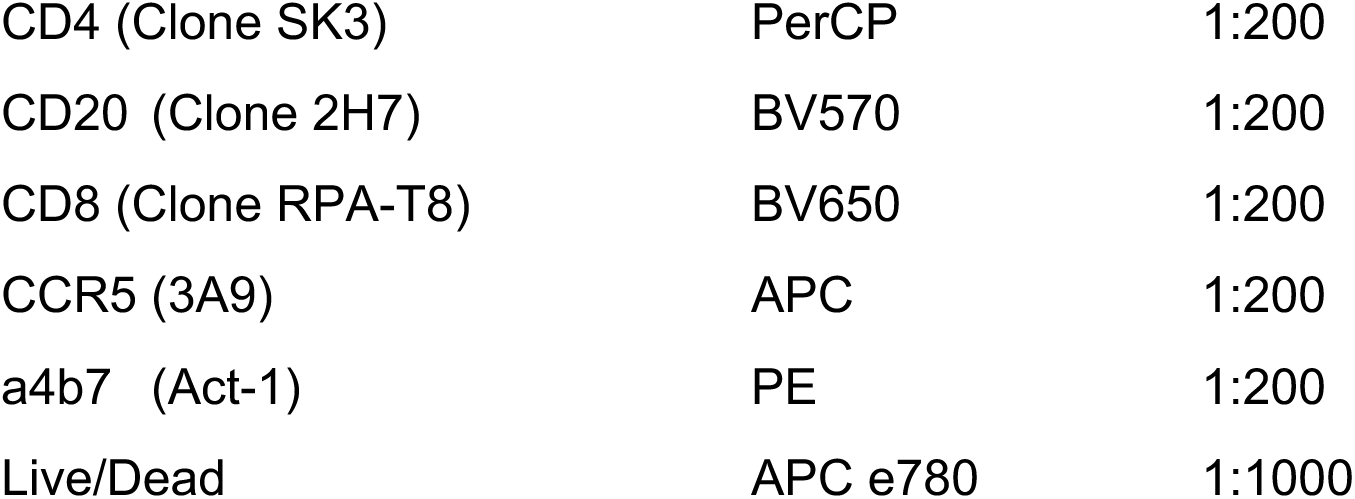

CD8 Intranuclear Panel:

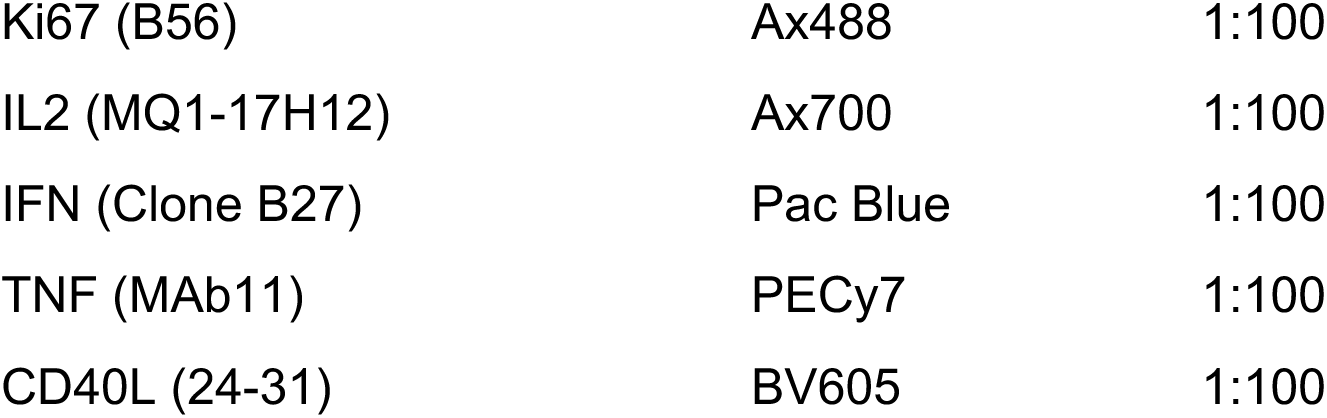

CD4 T Cell AIM Surface Marker Panel:

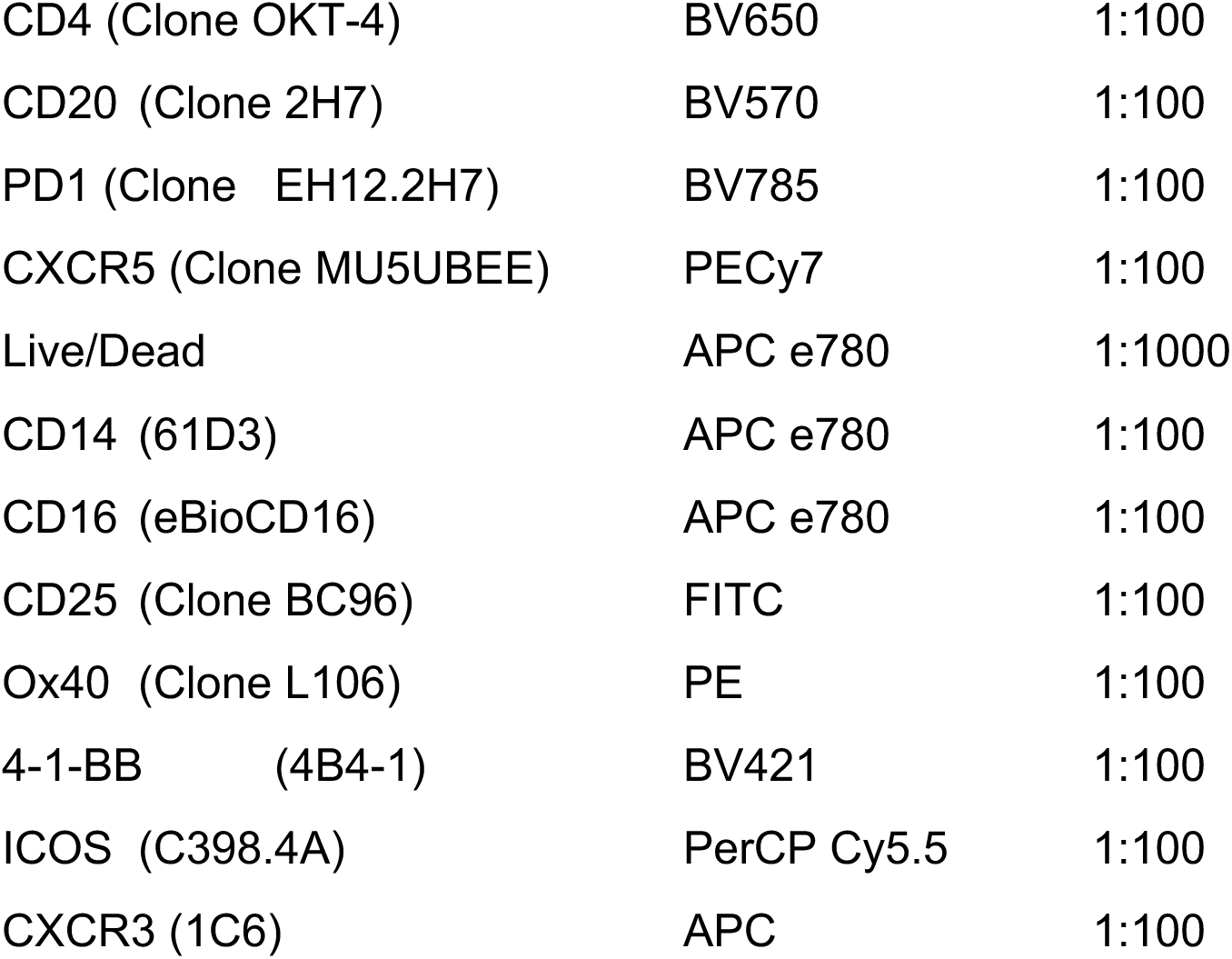

### Full length *env* viral sequencing

#### Long-read env sequencing

Samples were processed using the full-length Env sequencing protocol developed in (Laird Smith et al., 2016), but with modified primers and PCR conditions. Briefly, plasma samples were pelleted through a sucrose cushion to enrich for virions, RNA was extracted using the QIAamp Viral RNA Mini Kit (part no. 52906; Qiagen, Valencia, CA), and cDNA generated using the SuperScript III First Strand Synthesis System for RT-PCR (part no. 18080-051; Thermo Fisher, Fremont, CA), with oligo (dT) primers. SHIV *env* was amplified from this cDNA using the HIV *env* forward primer from (Laird Smith et al., 2016) Env-F: GAGCAGAAGACAGTGGCAATGA, and using a reverse primer designed for this SHIV: CCCTGATTGTATTTCTGTCCCTCAC, both purchased (de-salted) from Integrated DNA Technologies (San Diego, CA) and diluted to 20 pmol in 0.1X TE buffer before use. PCR was as in (Laird Smith et al., 2016), using the Advantage 2 PCR reaction mixture (Advantage 2 PCR Kit, catalog no. 639206; Clontech, Mountain View, CA), with the SA Buffer, but using 42 cycles of 15 sec denaturation at 95°C, 30 sec annealing at 64°C, and 3 min extension at 68°C. A QIAquick PCR Purification Kit (part no. 28106; Qiagen, Valencia, CA) was used to purify PCR products, and Pacific Biosciences library preparation was exactly as in (Laird Smith et al., 2016), but using the newer P6/C4 chemistry, and with a modified 0.025nM loading concentration, and a 6 hour movie time. The challenge stock was handled identically but was highly concentrated and thus only 23 PCR cycles were used during amplification.

#### PacBio env data processing

An updated version of the Full-Length Env Analyzer (Eren et al., 2017; Laird Smith et al., 2016) pipeline was used to process SIV PacBio reads. Briefly, PacBio’s CCS2 algorithm was used to reconstruct single molecule Circular Consensus Sequence (CCS) reads, outputting fastq files. These reads were filtered for length, quality, and for matching an Env reference database (here we included the known BG505.SHIV challenge sequence) with FLEA’s default parameter settings. FLEA’s error correction and data-summarizing approach was used, again with default parameters, collapsing near-identical reads and generating high-quality consensus sequences (HQCSs), along with HQCS frequencies, which are then codon aligned. These HQCS sequences are visualized in a web browser environment, allowing the exploration of immunotype frequencies, and displaying variants upon the leaf nodes of a maximum likelihood phylogeny. Variant frequencies in Figure S3A-B were computed from HQCS sequence frequencies.

### Complex preparation for negative-stain EM

Serum Fab preparation was carried out as previously described (Bianchi et al., 2018). In brief, after buffer exchanging into TBS, up to ~1 mg of total Fab was incubated overnight with 10-15 μg BG505 trimers at RT in ~50 μL total volume. Complexes were then purified via size exclusion chromatography (SEC) using Superose 6 Increase 10/300 column (GE Healthcare) in order to remove unbound Fab. The flow-through fractions containing the complexes were pooled and concentrated using 100 kDa cutoff centrifugal filters (EMD Millipore). The final trimer concentration was titrated to 0.04 mg/mL prior to application onto carbon-coated copper grids.

### Negative-stain EM

The SEC-purified complexes were applied to glow-discharged, carbon-coated 400-mesh copper grids, followed by pipetting 3 μl of 2% (w/v) uranyl formate stain and blotting, followed by application of another 3 μl of stain for 45–60 s, again followed by blotting. Stained grids were stored under ambient conditions until ready for imaging. Images were collected via Leginon software using a Tecnai T12 electron microscopes operated at 120 kV ×52,000 magnification. In all cases, the electron dose was 25 e−/A2. Particles were picked from the raw images using DoG Picker and placed into stacks using Appion software. 2D reference-free alignment was performed using iterative MSA/MRA. The particle stacks were then converted from IMAGIC to RELION-formatted MRC stacks and subjected to RELION 2.1 2D and 3D classification. A detailed protocol can be found in Bianchi et al., Immunity 2018.

## QUANTIFICATION AND STATISTICAL ANALYSIS

Infection probability per challenge event was modeled as depending on the BG505 N332 S375Y log10 ID_50_ nAb titer at the time of challenge using a modified logistic regression, where the maximum infection probability (where 0 < max < 1) was an additional parameter to be estimated by the model, rather than being fixed at 1 as in traditional logistic regression:

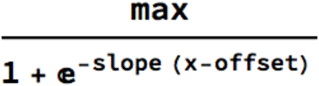

This adjustment is necessary because unimmunized animals with no serum nAb titers are not infected with 100% probability upon the first challenge, as a consequence of the chosen AID_75_ challenge dose. The infection event was assumed to be the challenge time point prior to the detection of viremia. Per-time point challenge outcomes were assumed to be conditionally independent of each other when conditioning on the corresponding BG505 N332 S375Y log10 ID_50_ nAb titer of the respective time point. We assumed weakly informative priors over the three model parameters, with slope~*Normal*(0,10), offset~*Normal*(0,10), and max~*Uniform*(0,1), and we used the Metropolis algorithm to draw 1 million samples from the posterior distribution. Chain mixing was rapid (see trace plots in Figure S5B), with effective sample sizes (ESSs) above 20,000 for all 3 parameters and for the log posterior probability. The posterior parameter distributions are visualized in Figure S5A. The calculated 5%, 50%, and 95% quantiles for each parameter were:

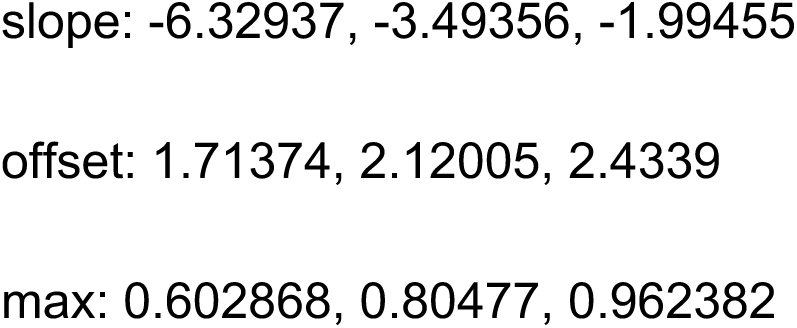

While under the prior distribution, P(slope < 0) = 0.5 and P(slope > 0) = 0.5, allowing equal prior probability of protective or sensitizing effects of neutralizing antibodies, the posterior probability of P(slope < 0) = 1 indicated the strongest possible evidence for decreasing infection probabilities given increasing ID_50_ nAb titers. Figure S5C shows 10,000 posterior sampled logistic curves, and the 5%, median, and 95% credible intervals for the infection probability computed from these, that were used to plot Figure 4C.

Graphpad Prism v7.0 was used for all standard statistical analyses. The significance of differences in neutralization and binding data between groups was calculated using unpaired, two-tailed Mann-Whitney U tests, correlations were calculated using Spearman correlation tests. Statistical parameters of all analyses are reported in the respective figure legends.

## KEY RESOURCES TABLE

